# Challenges in screening for *de novo* noncoding variants contributing to genetically complex phenotypes

**DOI:** 10.1101/2022.11.05.515231

**Authors:** Christopher P. Castro, Adam G. Diehl, Alan P. Boyle

**Affiliations:** Department of Human Genetics, University of Michigan, Ann Arbor, MI, 48109, USA; Department of Computational Medicine and Bioinformatics, University of Michigan, Ann Arbor, MI, 48109, USA

## Abstract

Understanding the genetic basis for complex, heterogeneous disorders, such as autism spectrum disorder (ASD), is a persistent challenge in human medicine. Owing to their phenotypic complexity, the genetic mechanisms underlying these disorders may be highly variable across individual patients. Furthermore, much of their heritability is unexplained by known regulatory or coding variants. Indeed, there is evidence that much of the causal genetic variation stems from rare and *de novo* variants arising from ongoing mutation. These variants occur mostly in noncoding regions, likely affecting regulatory processes for genes linked to the phenotype of interest. However, because there is no uniform code for assessing regulatory function, it is difficult to separate these mutations into likely functional and nonfunctional subsets. This makes finding associations between complex diseases and potentially causal *de novo* single-nucleotide variants (dnSNVs) a difficult task. To date, all but one published study in this area has failed to find any significant associations between dnSNVs from ASD patients and any class of known regulatory elements. We sought to identify the underlying reasons for this and present strategies for overcoming these challenges. We show that, contrary to previous claims, the main reason for failure to find robust statistical enrichments is not the number of families sampled, but the quality and relevance to ASD of the annotations used to prioritize dnSNVs, and the reliability of the set of dnSNVs itself. We present a list of recommendations for designing future studies of this sort that will help researchers avoid common pitfalls.

## Introduction

In human medicine, heterogenous phenotypes are frequently grouped into single disorders based solely on similarities in clinical presentation. Diagnostic criteria for these conditions usually consist of lists of symptoms, of which individual patients typically exhibit only a subset. As such, it is possible for two patients to display completely disparate sets of symptoms while still meeting the diagnostic criteria for said disorder. Identifying the genetic basis of these disorders is necessary for understanding their underlying mechanisms and developing effective treatments. However, the breadth of phenotypes presented by individuals sharing the same diagnosis reflects an underlying genetic basis that is at least as complex. Indeed, the combination of subjective diagnostic criteria and the likely polygenic basis of most diagnostic symptoms, makes it possible for individuals with the same diagnosis to have completely distinct sets of underlying mutations affecting entirely different pathways. Dissecting this variation is necessary to identify common themes in the etiology and underlying mechanisms of these disorders.

Among these disorders, Autism Spectrum Disorder (ASD) stands out as one of the most complex, making it a good case study for developing robust statistical methods to identify novel variants contributing to complex disorders. Such mutations are difficult to identify using traditional statistical methods owing to difficulty objectively grouping individual patients into cohorts and the fact that even patients with similar clinical presentation may not share the same underlying genetic and mechanistic basis. Here we use ASD to model complex, heterogeneous phenotypes, and test strategies for identifying *de novo* variants relevant to ASD given currently-available whole-genome-sequencing (WGS) datasets and functional genomic annotations.

ASD is a term used to describe a group of neurodevelopmental conditions often characterized by difficulties with social interaction, communication, and behavior. A genetic basis for ASD was first established through twin studies showing stronger concordance between monozygotic siblings compared to dizygotic siblings [1][2], [3]. Most estimates now place the heritability of ASD in the range of 50%-90% [4]–[7]. With technological improvements, several classes of genetic variations have been revealed to contribute to ASD etiology. This includes point mutations and structural variants (such as copy-number variants), pointing to a genetically heterogeneous background [8][9]. It is clear that many different types of genetic factors play a role in ASD, and it would benefit the research community to begin bridging the gaps in our understanding of the underlying genetic heterogeneity. As a whole, common variants contribute strongly to ASD. Individually, however, each of these common variants are expected to have small effects. This can be explained, in part, by the fact that variants associated with large, harmful effects are less likely than neutral variants to be maintained in a population. Conversely, uninherited variants that arise spontaneously (*de novo* variants) may carry a higher risk than inherited variants because they have not yet been acted upon and removed by natural selection. While the combined effects of common variants may contribute to a large portion of the heritability of autism [4], *de novo* mutations may potentially have larger individual impacts.

A significant role for protein-coding variants in ASD has been established [10]. Still, the coding variants that have been identified account for only a fraction of the overall heritability of ASD. There is evidence that the genetic background of ASD likely involves a combination of both coding and noncoding variation. Although effect sizes of mutations in coding regions may, on average, be greater because of their ability to directly affect amino acid sequences, mutations in noncoding regions may contribute to autistic phenotypes in an alternate way: by disrupting regulatory sequences [10]. Regulatory elements, such as promoters and enhancers, are responsible for controlling the precise time, location, and level of gene expression. Mutations disrupting regulatory elements may interfere with proper expression of developmental genes in the brain, for example, leading to phenotypes that are characteristic of ASD. With more than 98% of the human genome composed of noncoding DNA, those noncoding regions present a logical place to potentially uncover some of the missing heritability of autism. Indeed, most genome-wide association study signals map to noncoding regions, highlighting their importance. Therefore, investigation of noncoding variation has the potential to uncover novel ASD-associated variants.

Beyond the identification of noncoding variants, the challenge of their functional interpretation remains. Previous studies have investigated possible roles of *de novo* single-nucleotide variants (dnSNVs) in ASD by testing for enrichment of functional evidence among dnSNVs in probands versus siblings used as matched controls. However, while these studies uncovered significant associations with several coding categories, ascertaining the functional impact of noncoding dnSNVs is more difficult: whereas known properties of open reading frames, splice sites, and the biochemical properties of amino acids facilitate coding dnSNV prioritization, no such code exists to prioritize noncoding dnSNVs. Despite this difficulty, previous work has implied a modest contribution of *de novo* noncoding variation in autism, although coding regions exhibited the strongest associations and noncoding associations were not robust to multiple-testing correction [11]. The authors did not suggest that a noncoding association does not exist, rather, that the signal is not as strong as expected and that larger cohorts and careful attention to multiple-testing burden would be necessary to observe any true signal from *de novo* noncoding mutations. Indeed, other studies which have focused on noncoding mutations in ASD have also seen significant enrichment disappear once multiple-testing corrections have been applied [12]. Nonetheless, these studies establish that these rare mutations do play a role in ASD, underscoring the importance of improving methods with which to study them.

In recent years, we have learned certain features associated with active regulatory sequence, including epigenetic marks, sequence motifs for various TFs, chromatin accessibility, etc. Although we know that regulatory sequences are associated with characteristic combinations of functional annotations (H3K4me1 and open chromatin for enhancers, e.g.), our ability to predict the regulatory function of a locus based on overlapping annotations remains limited. Despite an abundance of publicly-available functional genomics datasets across many tissue and cell types, a uniform functional code for regulatory sequences remains elusive. In the absence of such a code, variants must be prioritized based solely on their overlap with various combinations of functional annotations thought to be important for regulatory function. Several approaches for screening and combining annotations have been used to prioritize variants in the ASD literature with varying degrees of success.

Most studies to date have leveraged the large number of public datasets by combining individual functional annotations into functional categories meant to reflect the regulatory potential of individual variants. A statistical test for enrichment must then be run on each of these functional categories to identify if an excess of dnSNVs exists in probands, which may signal an association with development of ASD. Since it is not known in advance which categories will be informative, this strategy requires many individual tests to comprehensively screen for associations. Unfortunately, as the number of annotations increases, the number of individual tests increases exponentially, invoking a substantial multiple-testing burden and potentially reducing statistical power by orders of magnitude. As a result, very few studies have been able to identify robust statistical enrichment of dnSNVs within any functional category in ASD affected individuals. In nearly all of these studies, the solution presented for this problem is to increase the number of families until sufficient power is achieved.

We wanted to explore the implications and practicality of this solution in comparison to newer methods of variant prioritization, in hopes of identifying the optimal solution given current technology. To this end, we analyzed whole-genome sequencing data from 1,917 quad families in the Simons Foundation Autism Research Initiative (SFARI) Simons Simplex Collection (SSC) cohort. These families represent a total of 7,668 individuals: 1,917 affected children and 5,751 unaffected parents and siblings. This family structure is ideal for identifying *de novo* variants and provides us with a natural control group in the unaffected siblings. Using a set of newly-identified dnSNVs from this cohort, we quantified the ability of selected combinations of individual annotations, and scores from published variant-prioritization methods, to detect differential associations with dnSNVs. For each of these functional categories, we used 80% power curves to predict the optimal sample size, at which 80% of tests are expected to be significant. We further evaluated the relative effect size necessary to find a significant result given the current sample size in each functional category, expressed as the difference in dnSNV burden in probands versus siblings using 80% power curves. Finally, we used simulations to compare the actual effect sizes observed in probands and siblings to random expectations to assess whether a significant test at any sample size is likely to reflect biologically-meaningful effects on ASD risk. Based on these experiments, we can draw several conclusions about current limitations in our ability to confidently identify dnSNVs contributing to ASD, and propose strategies to improve future studies on the impact of dnSNVs on ASD and other complex genetic disorders.

## Materials and Methods

### Identification and filtering of de novo single-nucleotide variants

In order to identify candidate *de novo* SNVs while minimizing false positives, we implemented a pipeline to improve the quality of genotype and variant calls before applying stringent filters. We first performed genotype refinement using GATK’s [13] CalculateGenotypePosteriors tool and required all variants to pass the Variant Quality Score Recalibration (VQSR) filter, using a sensitivity threshold of 99.8%. We then filtered the variant set down to bi-allelic SNVs and tagged potential *de novo* mutations if a variant was present in a child and not any of the other family members, with the requirement that all four family members have GQ >= 20, DP >= 10, and the more stringent of AC < 4 or AF < 0.1%. This same process was followed separately both for probands and unaffected siblings to identify *de novo* SNVs for each group. We further filtered mutations down to exclude those appearing in more than one individual. Because the CalculateGenotypePosteriors tool was only designed to handle trios, we created separate PED files for probands and unaffected siblings, based on the PLINK pedigree file format. We referenced a file mapping SSC individual IDs to SSC family IDs, provided as a resource by the SSC, to generate these pedigree files.

We annotated all SNVs with their minor allele frequencies (MAF) using the Genome Aggregation Database (gnomAD) v2 [14]. Following annotation, we filtered the variant set down to those with a MAF less than .001. Variants for which a MAF was not available were also retained. We then used the Picard LiftoverVCF tool [15] to liftover variants from the hg38 build to hg19 since not all computational tools we used supported the hg38 build at the time of analysis.

Some families for which data was collected were initially enrolled in the SSC as simplex families but were later discovered to be multiplex and flagged as such by SFARI. These families were excluded from our analyses. Families were also excluded if they were part of the Simons Ancillary Collection or the Simons Twins Collection. To ensure that our identified mutations were true SNVs, variants overlapping with low-complexity regions were filtered out using UCSC’s RepeatMasker [16] and TRF [17] reference files. Additionally, the Mills and 1000g gold standard set was used as a reference for filtering out mutations overlapping INDELS [18].

### Annotation of coding dnSNVs

We defined coding mutations predicted to be damaging using Variant Effect Predictor (VEP) [19] annotations. For generating predictions, we used the “-most_severe” tag in order to ensure that each variant was assigned just one “consequence” instead of taking every possible transcript into account for that variant. Variants were considered to be “high-impact” dependent on their predicted consequence, based on the VEP variant consequence table from Ensembl [20]. Additionally, we considered variants to be predicted loss-of-function if they were annotated by SIFT [21] as “deleterious” and by Polyphen [22] as “probably damaging”. To identify genes that may be more susceptible to being affected by mutations, we annotated mutations with a score developed by the Exome Aggregation Consortium (ExAC) called pLI, which indicates a gene’s probability of being intolerant to loss-of-function (LoF) mutations [23]. A high pLI score implies a gene is LoF intolerant and we annotated genes with pLI>= 0.9 as extremely LoF intolerant. We downloaded pLI scores from https://gnomad.broadinstitute.org/downloads under “Gene Constraint Scores” [14].

### Annotation of fetal brain enhancer and promoter regions

To identify enhancer regions, we used combined male and female fetal brain DNase-seq and ChromHMM data from the Roadmap Epigenomics Project [24]. Using core 15-state ChromHMM models we identified regions containing histone marks corresponding to the ChromHMM states “Genic Enhancers”, “Enhancers”, and “Bivalent Enhancer.” We called those regions as enhancers if they also overlapped with DNase peaks. We used a similar process to identify promoter regions, focusing instead on the ChromHMM states “Active TSS”, “Flanking Active TSS”, and “Bivalent TSS”, before determining which of those regions overlapped with DNase peaks. We also annotated promoters through an alternate method using the GENCODE [25] release 19 gene annotation file. Using all protein-coding genes, we defined promoters as the region within 1,500bp upstream of the TSS of the respective gene.

### Annotation with functional scoring tools

We generated TURF generic and brain-specific scores as described in Dong and Boyle [26] with the tool available at https://github.com/Boyle-Lab/RegulomeDB-TURF. We generated Disease Impact Scores following the instructions for making predictions from the DNA model, provided at https://hb.flatironinstitute.org/asdbrowser/about.

### Other annotations

We obtained a “rank” for dnSNVs using the original RegulomeDB scoring system, with RegulomeDB v2.0. Ranks can be obtained from https://regulomedb.org/regulome-search. We considered mutations to have potentially disruptive regulatory effects if they received scores of 2 or 3 (note that a score of 1 is not possible in *de novo* mutations because it requires an eQTL annotation). For chromatin interaction analyses we used promoter-capture Hi-C data generated by Song et al. [27]. We applied those maps to our data to identify any potential contacts between dnSNVs and gene promoters. If such a contact was present, we assigned that mutation to the gene corresponding to that promoter. We annotated variants with CADD v1.4 [28] using offline scoring scripts, following instructions at https://github.com/kircherlab/CADD-scripts/ for the GRCh37 genome build. For identifying evolutionarily conserved elements we scored dnSNVs using the 46-way placental alignment phastCons [29] track from UCSC’s Genome Browser [30]. We obtained a list of genes, along with rankings based on strength of evidence of their association with ASD, from the SFARI Gene database at https://gene.sfari.org/database/human-gene/. We generated a list of genes that were found to be preferentially-expressed in brain tissue using data from A.B. Wells et al. [31].

### Enrichment testing procedures

For each of the annotations tested, we compiled contingency tables for Fisher’s exact test (FET) by counting the number of proband and sibling dnSNVs overlapping genomic regions falling within the category and those falling outside the category. Individual FETs were performed for each annotation category using the fisher.test method in R, with alternative=“greater” [32]. When multiple tests were performed, multiple-testing adjustments were made using the FDR method in R [33], [34]. For TURF generic, TURF brain-specific, and DIS scores, FET contingency tables were constructed by counting proband and sibling dnSNVs scoring in the top 5% of scores for each category. These were subjected to FET as described above. Wilcoxon rank-sum tests were also performed on non thresholded data from these categories to compare average score rankings between proband and sibling cohorts. For all tests, p <= 0.05 was used as the significance threshold.

### Power analysis procedures

We performed the power analyses in R using the ‘pwr’ package. To calculate the power for each annotation category across varying sample sizes we implemented the two-proportion test within the ‘pwr’ package. Proband and sibling proportions were defined as the proportion of dnSNVs within a particular annotation category compared to the total number of dnSNVs in probands and siblings, respectively. We ran each calculation at a significance level of 0.05, and with the alternative hypothesis being “greater.”

### Reverse power analysis procedures

We produced “reverse” power curves, where we held the sample size fixed and plotted power over a range of differential overlapping proportions of proband and sibling dnSNVs for each annotation category in order to assess how much additional information each would need to convey in order to reach 80% power with 1,917 quad families. The same methods and thresholds described in the “Power analysis procedures” were then used to plot power curves with the ‘pwr’ R package.

### Comparison to random permutations

In order to assess whether observed counts for each of our annotation categories significantly deviate from random expectations, we performed a permutation analysis using the same input data used for the FETs. Data were randomized by shuffling the “proband” and “sibling” labels across all dnSNVs, for 10,000 permutations. For each permutation, we stored the number of proband and sibling dnSNVs overlapping each annotation category in the permuted data. Counts were used to generate an eCDF for each annotation category, and the mean of this distribution was used as the expected count for proband and sibling. We then calculated z-scores to quantify the deviation between the observed proband and sibling counts and the expected randomized mean based on the standard deviation of the eCDF. Z-scores of 2.0 or greater were considered significant evidence for departure from random expectation.

### Comparison of dnSNV datasets across studies

For comparisons between studies, we obtained the publicly-available dnSNV datasets from each of the other groups we included in our analyses. As with our own set, we restricted the variants to autosomal dnSNVs. We generated the UpSet plot using Intervene [35] and UpSetR [36]. For direct comparison between our variant set and that of Zhou et al., we used BEDTools [36], [37] to obtain the intersection, union, and disjoint sets.

## Results

### De novo SNV calls show substantial overlap with previous studies

From the SSC, we identified a median count of 70 autosomal *de novo* single-nucleotide variants per proband (134,969 total autosomal proband dnSNVs). This is consistent with the estimated mutation rate in the general human population [38], [39]. Compared to the median of 68 autosomal *de novo* mutations we identified in the unaffected siblings (131,896 total autosomal sibling dnSNVs), we did not observe a statistically significant difference in dnSNV count between the proband and sibling groups (Fig. 1A).

**Figure 1.**
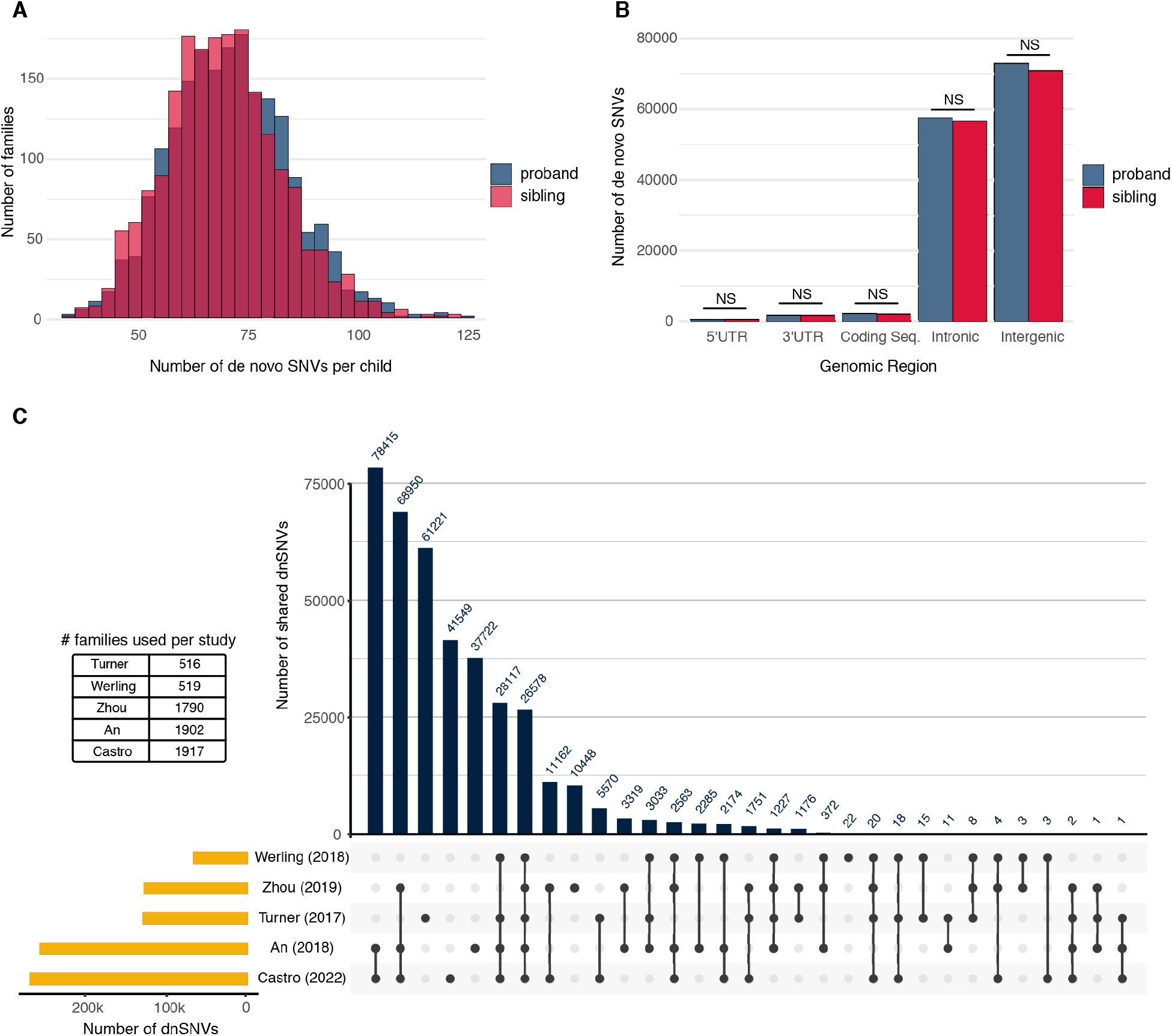
Distribution of *de novo* single-nucleotide variants from whole-genome sequencing of the Simon’s Simplex Collection. **(A)** Distribution of *de novo* SNVs (dnSNVs) across 1,917 families from the SSC. Probands (blue) have a median of 70 dnSNVs per child while the median for unaffected siblings (light pink) is 68 dnSNVs. The darker pink bars indicate overlap in counts between probands and siblings. The difference in counts between the two groups is not statistically significant **(B)** Distribution of dnSNVs by genomic region. Approximately 98% of all dnSNVs identified and used in this study land in intronic or intergenic regions. The number of dnSNVs in each category is not significantly different between probands and siblings. **(C)** Total number of dnSNVs dentified across different individual studies (gold bars), all using the Simons Simplex Collection cohort data. Blue vertical bars indicate the number of variants identified by more than one study (solid black points connected by black line) or variants only identified by a single group (solid black point). Although all families are part of the same cohort, the number of families utilized by each study varies, shown in the table.

We annotated and categorized the dnSNVs into the following genomic regions: 3’UTR, 5’UTR, intergenic, intronic, and coding. Roughly 98% of dnSNVs overlap noncoding regions of the genome, which falls in line with coding regions comprising ∼1.5% of the human genome. When taking all dnSNVs into account, within each genomic region, there is not a statistically significant difference in counts between probands and siblings (Fig 1B).

We compared our list of identified dnSNVs to those from four other published studies which also used the SCC data (Fig. 1C). Approximately 84% of the dnSNVs we identified have also been identified by at least one of these four groups [11], [12], [40], [41]. While all compared groups used data from the same cohort (SSC), the number of cohort families included in each group’s respective analyses varied. The largest number of variants shared between studies came from the intersection of our study (1,917 families included in analyses) with that of An et al. (1,902 families). The next highest overlap in variants came from the intersection between three studies: our own, An et al. and Zhou et al. (1,790 families). As would be expected, there was a smaller amount of overlap with the studies which included fewer families on account of data availability at the time of their publication (Turner et al. – 516 families, Werling et al. – 519 families).

### De novo coding variants show significant association with ASD

In order to validate our dnSNV calling and enrichment testing strategies, we wanted to first show our ability to recover enrichments of high-impact coding mutations, which have been previously shown to be significantly associated with ASD [10], within proband dnSNVs. We prioritized dnSNVs by annotating them using Variant Effect Predictor (VEP) [19], SIFT [21], and PolyPhen [22]. Mutations were classified as predicted loss-of-function coding mutations (LOFCMs) if they were annotated as “high-impact” by VEP, or as both “deleterious” by SIFT and “probably damaging” by Polyphen. We used Fisher’s Exact Test (FET) to weigh the evidence for a statistically significant excess of predicted LOFCMs in probands compared to siblings.

Consistent with our expectations, we observed 604 proband LOFCMs compared to 467 sibling LOFCMs, representing a statistically-significant enrichment (FET, FDR-adjusted p=0.002)(Fig. 2). In all, over 90% of VEP high-impact variants were stop-gain mutations, which result in premature stop codons and, in turn, truncated transcripts. These stop-gain mutations were enriched even more strongly than all LOFCMs, with nearly twice as many found in probands compared to siblings (120 vs 62 mutations; FET, FDR-adjusted p=0.001). Taken together, these observations show we are able to recover known enrichments within our dnSNV dataset.

**Figure 2.**
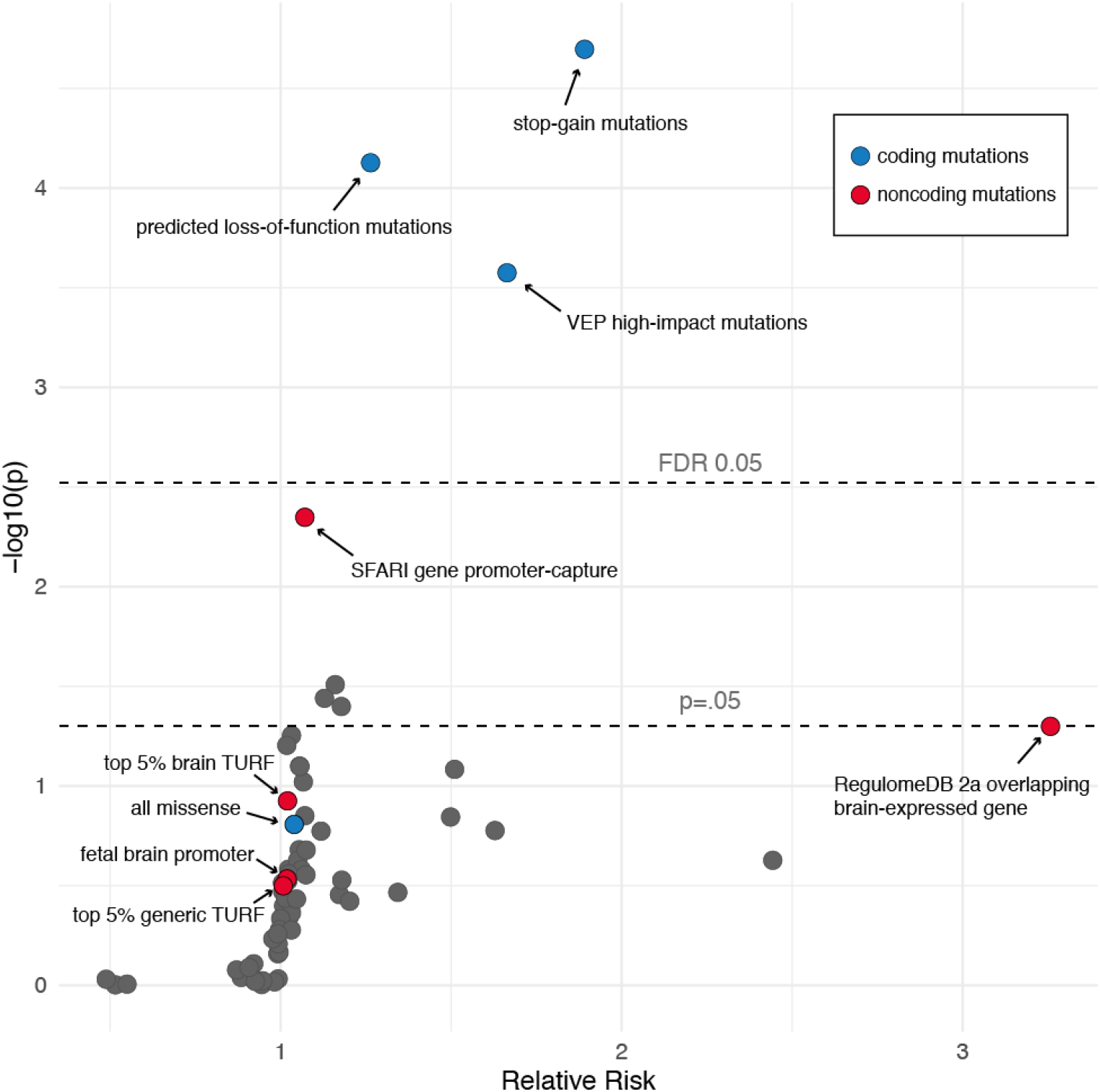
Relative risk of proband dnSNVs across 65 annotation categories, including combinations of different annotations. A relative risk >1 represents enrichment of dnSNVs in the proband group. The only categories that remain significant after multiple-testing correction are related to coding-region annotations.

### Proband dnSNVs are not enriched for predicted regulatory variants

We next wanted to extend our enrichment testing strategy to noncoding mutations. Specifically, we were interested in determining if we could detect an enrichment of proband mutations with the potential to affect regulatory function. We wanted to know whether a more comprehensive set of dnSNVs coupled with our FET screening strategy would yield any statistically significant enrichments in noncoding regulatory annotation categories.

This approach relies on our ability to predict how likely noncoding variants are to affect regulatory function. Most published studies have done so based on overlap with genomic annotations commonly associated with cis-regulatory regions. Annotations are derived from the Encyclopedia of DNA Elements (ENCODE) project and include, open chromatin (DNase-seq and DNase footprinting), and transcription factor (TF) binding sites (ChIP-seq), among others. These are commonly combined into annotation categories; either exhaustively, by selecting a subset manually, or by using computational methods. However, more recent studies have turned to machine learning to identify the relationships between functional annotations and ASD. The distinct advantage of this approach is that it may reduce or obviate the need for multiple-testing corrections. In order to compare these strategies, we selected a combination of annotation categories and prioritization scores and proceeded with enrichment tests to determine if any significant associations could be recovered.

As representative manually-curated annotation categories, we combined annotations from the ENCODE database for ChromHMM and DNase-Seq, enhancer and promoter predictions from the Roadmap Epigenomics Project [24], chromatin interaction data from Song et al. [27], and brain-specific gene expression data from [31]. These were used to isolate sets of likely promoter and enhancer regions targeting genes expressed in the brain. Given known features of ASD etiology, these two genomic compartments seemed likely to harbor an enrichment of dnSNVs affecting relevant regulatory functions. However, when considering the number of mutations overlapping enhancers we saw no significant difference between the proband and sibling groups (3114 vs 3053, respectively; p=0.79, FET). Similarly, there was no enrichment of mutations within promoters (1814 vs 1750; p=0.5, FET).

As representative variant prioritization methods, we selected Tissue-specific Unified Regulatory Features (TURF), a probabilistic scoring model that prioritizes in a tissue-specific manner, which replaced the original categorical scores in the current release of RegulomeDB [42]. For noncoding variants, we scored each dnSNV with two different TURF prediction scores; one score was based on a previous implementation of TURF which scores variants in a generic context (independent of tissue) [43]. The second score was generated by the current implementation of TURF, in which functional variants are predicted in a tissue-specific context [26]. We calculated these scores based on functional evidence specifically from brain tissue, which we could reasonably expect to be more relevant to ASD. For both generic and brain-specific TURF scores, FET contingency tables were constructed based on overlap with positions scoring in the top 5% of annotated sites. In both cases, tests were not significant after multiple-testing correction, similar to previous studies (generic TURF: FET, FDR-adjusted p=0.6; brain-specific TURF: FET, FDR-adjusted p=0.5)(Fig. 2). Since TURF scores are numeric and continuous, we retested for enrichment using Wilcoxon Rank-Sum tests, which also failed to reach significance (generic TURF: Wilcoxon Rank-Sum, p=0.83; brain-specific TURF: Wilcoxon Rank-Sum p=0.89).

### Tissue- and disease-specific annotations are more informative than tissue-agnostic annotations

Given the lack of enrichments observed for any annotation categories or prioritization scores, we wanted to further explore how informative these annotations actually are. Since most previous studies failing to show significant enrichments have cited insufficient sample size as their primary limitation, we first asked what sample size would be necessary to achieve 80% power in our statistical tests. To do so, we plotted power curves for three noncoding annotation categories and prioritization scores: TURF generic, TURF brain-specific, and Fetal Brain Promoters. For comparison, we included two coding annotation categories: missense variants, regardless of severity of impact, and high-impact variants, which are likely to lead to loss-of-function.

Considering a desired power threshold of at least 80%, our analyses revealed that we are underpowered to detect enrichment of dnSNVs in probands for any of the noncoding categories given our sample size of 1,917 quad families (Fig. 3A). As a baseline, we estimated a power of ∼97% when testing for enrichment of high-impact coding dnSNVs using the same sample size. However, it is notable that the missense coding category yielded only ∼27% power at the current sample size, slightly below the highest-performing noncoding category: TURF brain-specific scores (∼32% power). TURF brain-specific scores were estimated to reach 80% power at a sample size of ∼10,000 families.

**Figure 3.**
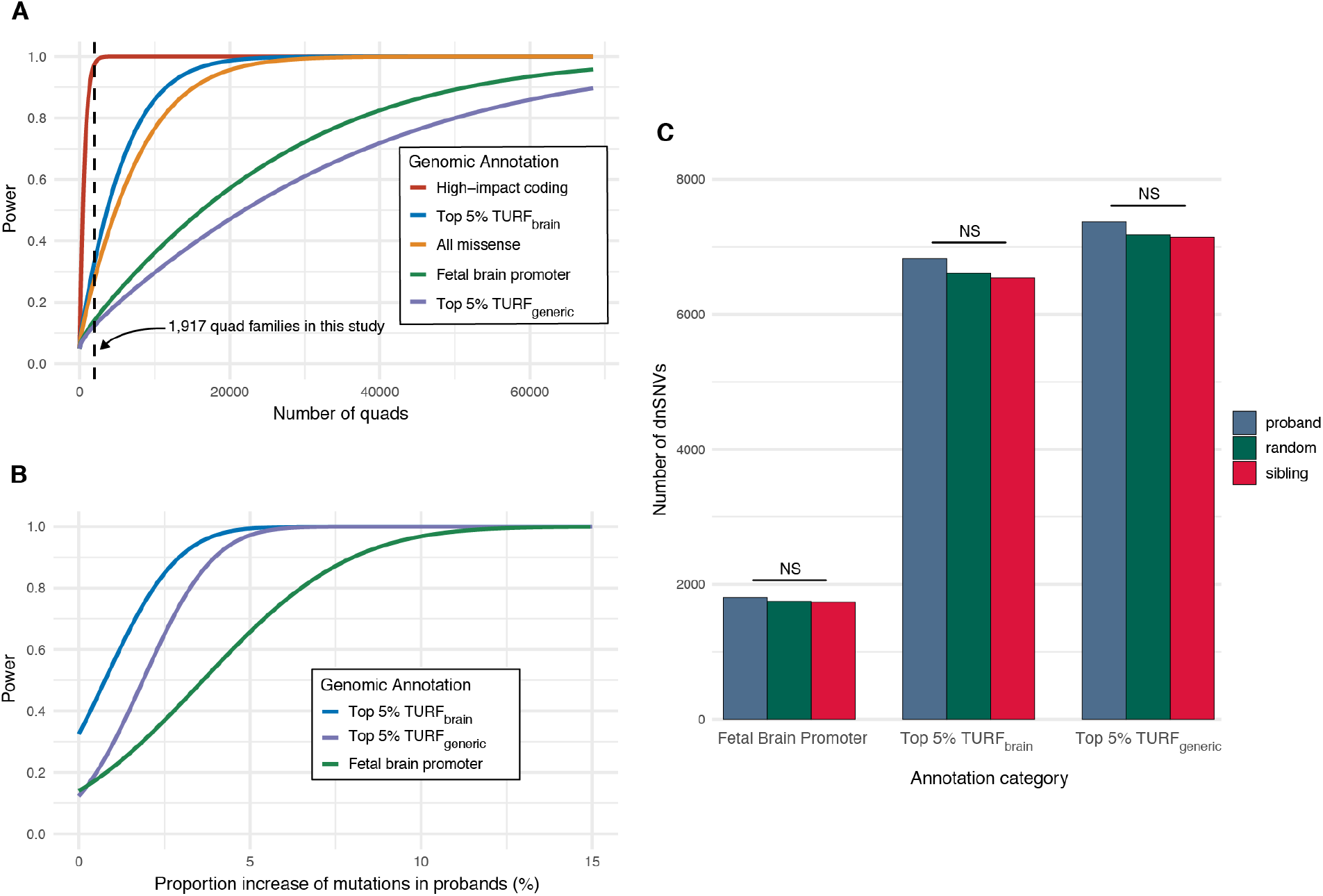
**A)** Power analysis for detecting proband enrichment of different categories of *de novo* SNVs. The black dashed line indicates our current sample size (1,917 quad families). We have estimated the sample sizes necessary in order to detect association of dnSNVs with ASD. We estimate a power of ∼97% when testing for enrichment of high-impact coding dnSNVs in probands at our current sample size. The missense coding category yields a power of 27%. Brain-specific TURF scores (32% power) would require ∼10,000 more families to achieve 80% power. Over 50,000 families would be necessary for generic TURF scores (12%) to reach that same 80% threshold. The fetal brain promoter category slightly outperforms the generic TURF scores at 13%. **B)** Generic TURF starting power = 0.12, achieves 80% at ∼3.2% increase (∼240 additional variants, 7,370 observed). Brain TURF starting power = 0.32, achieves 80% at ∼2.2% increase (∼150 additional variants, 6,828 observed). Fetal brain starting power = 0.14, achieves 80% at ∼6.5% increase (∼117 additional variants, 1,806 observed). **C)** Observed counts of proband (blue bars) and sibling (red bars) *de novo* SNVs prioritized with three different noncoding annotations. We observed no significant difference between random counts (green bars) and counts in probands or siblings (Z-scores: fetal brain promoters = 0.53, TURF generic = 0.48, TURF brain = 1.16, permutation tests).

We note that if we were to instead use generic (non-brain-specific) TURF scores for prioritization, over 50,000 families would be necessary to achieve 80% power, highlighting the potentially profound effects of the choice of training strategy for variant prioritization. In particular, we see that including data directly relevant to the tissue and developmental stage under study offers a significant improvement for this application. This is underscored by the observation that generic TURF scores (∼12% power) underperformed the manually-curated fetal brain promoter category (∼13% power) even though generic TURF scores incorporate more annotations to generate their prioritization scores. Even so, an estimated 37k families would be required to reach 80% power for detecting enrichment of fetal-brain promoter mutations, showing that neither of these annotation categories are particularly informative in terms of ASD risk.

### Improving annotation quality has more impact on empirical power than increasing sample size

We wanted to resolve the question of whether a significant test result based on a larger sample size would actually reflect a meaningful association between any of these annotation categories and ASD. In order to further dissect the strength of associations between proband dnSNVs and ASD, we assessed statistical power for each annotation category at a fixed sample size over a range of effect sizes. We defined the effect size for an annotation category as the difference between the fractions of proband and sibling dnSNVs within a given annotation category. The magnitude of this difference is indicative of the strength of the association between an annotation and ASD. A highly informative annotation category would be expected to associate mostly with proband dnSNVs and only rarely with sibling dnSNVs, so “overlapping” counts in the FET contingency table would be highly skewed toward the proband column. The effect size, therefore, would be relatively large. By contrast, an uninformative classifier would associate randomly with proband and sibling dnSNVs; i.e. the observed proportions of proband and sibling dnSNVs overlapping the annotation category would be equal. Thus, the observed effect size would be very small. We wanted to assess how informative our annotation categories actually are in differentiating between proband and sibling cohorts. More precisely, we wanted to ask, if we could improve an annotation to make it more informative, how much more information must it convey to achieve 80% power at the current sample size? We interpreted this as an ersatz quality metric for each annotation category.

We evaluated how much we would need to inflate the effect size of each annotation category by plotting curves relating statistical power to the proportion of excess information in probands relative to siblings in FETs. In this framework, zero effect size is observed when both proband and sibling have equal proportions of dnSNVs overlapping a given annotation category. We then emulate the consequences of increasing the effect size by increasing the proband overlapping proportion while holding the sibling proportion fixed, thus artificially increasing the effect size of the annotation category. By recalculating empirical power thusly over a range of effect sizes, we can plot curves showing the necessary effect size increase to reach 80% power at the current sample size (Figure 3B).

Consistent with our power analysis results, brain-specific TURF scores required the smallest increase in effect size to reach 80% power, approximately 2.2%, or roughly 150 more than the 6,828 we actually observed. This makes them somewhat more informative than generic TURF scores, which would require a 3.2% increase of dnSNVs overlapping the top 5% of scores, or 240 more dnSNVs than the 7,370 actually observed. However, both variant prioritization methods performed substantially better than the manually-curated fetal brain promoter category, which would require a 6.5% increase in information to achieve 80% power at the present sample size, or 117 variants in addition to the 1,806 observed.

### Comparison of current annotations to random permutations

Given the results of both the power and effect size analyses, we can conclude that the most-informative noncoding annotation category among those tested were TURF brain-specific scores. However, even though only a modest increase in either sample size or effect size was necessary to get to 80% power, the actual enrichment test still failed to reach significance. All others performed significantly worse: extreme increases in either sample or effect sizes would be necessary to reliably detect enrichments. This led us to question whether these annotation categories actually conveyed significantly more information than random expectations.

For each of the three noncoding annotation categories, we modeled the expected random overlap of proband and sibling dnSNVs with the given annotation category by randomly shuffling “proband” and “sibling” labels across all dnSNVs for 10,000 permutations. The mean number of overlapping proband and sibling dsSNVs across permutations were then compared to the applicable observed counts (Fig 3C). Z-scores were calculated to quantify the degree of departure between the observed count and its random expectation. Z-scores exceeding 2.0 were considered significant. However, the observed Z-scores for all categories were well under this threshold. Notably, TURF brain-specific scores, which performed the best in our other tests, was the only annotation category with a Z-score exceeding 1 (∼1.16). By comparison, the high-impact coding category produced a Z-score of ∼3.57 using these methods. Therefore, we can conclude that even the best of our annotation categories are relatively uninformative in regards to ASD risk or etiology.

### dnSNV calls show variable quality across studies

We have seen the limitations of sample size and variant effect size across several studies conducted by other research groups who have used the same raw data from the Simons Simplex Collection to study noncoding mutations in ASD. Interestingly, while all these studies are subject to the same limitations, their results appear to vary substantially in terms of the specific associations found, with little reproducibility between groups, even when testing methods were similar.

To date, only one study has reported a robust statistical enrichment in a noncoding category [41]. The authors use a disease impact score (DIS) to prioritize dnSNVs relative to their impact on brain disease. This method is an extension of DeepSea [44], a machine-learning model that assigns functional scores to individual variants based on overlap with functional annotations from ENCODE and other sources. DIS extends this model by way of training on a curated set of known features related to brain disease phenotypes. The authors found significant evidence for higher DIS scores in probands compared to siblings (p=.009, one-sided Wilcoxon rank sum test).

We wanted to know whether DIS scores would yield a significant enrichment within our data as well, so we tested our own set of dnSNVs but found no enrichment using either FET (p=0.214, one-sided FET) or Wilcoxon Rank-sum tests (p=.071, one-sided Wilcoxon rank-sum test). This prompted us to investigate the effect of the specific set of dnSNVs on test results. We investigated this by testing for association of DIS scores within the union, intersection, and disjoint fractions of our dnSNVs and those from Zhou et al. For each of these fractions, we repeated the Wilcoxon rank-sum test as before, making note of whether a significant difference was apparent.

The only significant result we observed was for the intersection of both datasets (p < 0.0026, one-sided Wilcoxon rank-sum test), which actually showed stronger evidence for enrichment than Zhou et al. originally reported (p=0.009, one-sided Wilcoxon rank-sum test). By contrast, neither the union (p=.083, one-sided Wilcoxon rank-sum) nor disjoint datasets produced a significant result, with the lowest performance observed for the disjoint sets (This study: p=0.99; Zhou et al.: p=0.73). This suggests that the quality of dnSNV calls varies, with the highest-quality calls also being the most reproducible.

We further tested this hypothesis by intersecting our variants with those from the previously-mentioned four other groups who have used their own methods to identify dnSNVs in the SSC cohort (Fig 1c). We filtered down our set of dnSNVs by keeping only those which appeared in at least two of these other groups’ published sets. Once again, performing the Wilcoxon rank sum test on the intersection set of dnSNVs revealed a significant difference in DIS between probands and siblings (p = .02837), albeit at lower confidence compared to the intersection between our dnSNVs and those from Zhou et al. We speculate that this decrease results from the decrease in overall sample size incurred due to the smaller absolute size of the dnSNV datasets in other studies.

Altogether, these results suggest that intersecting variants across sets of calls leads to an overall increase in quality among the dnSNV calls. We postulate that the disjoint sets of variants from across studies are enriched for false-positive variant calls, which may arise due to sequencing errors, genotyping errors, or other unknown sources. If not filtered out, these false-positive dnSNVs may dilute the signal from true dnSNVs sufficiently to prevent a significant test result even when an annotation is genuinely associated with ASD.

## Discussion

Prior to this analysis, several groups have used data from the SSC to seek associations between noncoding dnSNVs and ASD, with all but one yielding no statistically significant enrichments. Our results on three different noncoding annotation categories were consistent with their results in that we found no significant associations. Our testing methods reproduced previously-demonstrated enrichments of proband dnSNVs within high-impact coding annotation categories. However, it was not immediately clear if we observed no noncoding enrichments due to insufficient sample size, as suggested by the authors of most previous studies, or the inability of the annotations we chose to reliably differentiate between regulatory and neutral noncoding variants.

In order to systematically investigate these possibilities, we set out to objectively evaluate how informative current annotations are in regards to differences in functional effects of dnSNVs in probands vs siblings. This allowed us to compare different strategies in terms of their effects on our ability to find potentially-revealing genomic associations. Specifically, we explored the impact of sample size, choice of annotations/variant prioritization methods, and choice of variant sets, on our power to detect associations between *de novo* noncoding genetic variation and ASD.

Based on observed dnSNV counts in probands and siblings, our power analysis suggests at least 10,000 quad families would be necessary to achieve a power of at least 80% to detect an association between our best-performing noncoding annotations category and ASD. Although autism cohorts are constantly growing, this is approximately five times as many quad families than are currently available in the SSC. More importantly, though, the necessity of such a large number of families suggests a very small effect size, begging the question of whether such effects are meaningful. We show evidence that, in fact, current annotations are only slightly more informative than random expectations. Therefore, the strategy of increasing sample size alone is likely to lead to erroneous conclusions.

What we see from this is that it is not only the sample size that is limiting our ability to detect the effects of the dnSNVs, but also their effect sizes. This suggests that certain annotations categories may not sufficiently capture meaningful differences between probands and siblings: i.e., the annotations used are unable to reliably differentiate between noncoding dnSNVs that are neutral (without regulatory effects) and those capable of disrupting regulatory function. This may be amplified particularly in phenotypically heterogeneous disorders such as ASD. Therefore, an insignificant test result may not actually reflect lack of signal, but that the signal has been attenuated by high-scoring dnSNVs that actually lack functional significance. This effect may be particularly problematic if we rely solely on our intuition when choosing annotation categories. This is demonstrated by the poor performance of the fetal-brain-promoter category in this study; even though it is based on relevant, tissue-specific annotations, it still fails to produce a significant association. Thus, the most important choice when designing an experiment is likely which annotation category/ies and/or prioritization score(s) to use, keeping in mind the need to minimize multiple-testing burden. Werling et al. illustrated the importance of multiple-testing burden in a study that included a comprehensive set of >13k annotation categories, among which no significant associations were found after correcting for multiple-tests [11].

Accordingly, it is likely that improving annotations and prioritization scores, particularly in relation to their relevance to the specific tissue/disorder under study, is more likely to yield meaningful performance gains than increasing the number of available families for study. For example, we note that brain-specific TURF scores performed significantly better than generic TURF scores, highlighting the importance of using tissue-specific annotations when possible. We estimate nearly five times as many families would be necessary to achieve a power of 80% when using the generic TURF scores compared to tissue-specific TURF scores. Furthermore, DIS scores, which are specifically trained on disease-related features, outperformed TURF brain-specific scores even though the underlying training feature sets share substantial overlaps.

A clear limitation when investigating the impact of noncoding mutations in ASD is there are a great deal of ways with which we can choose to annotate and prioritize variants. In theory, we isolate the variants with evidence of being functionally relevant so that we can then use that subset of variants to test for genotype-phenotype associations. However, depending on the choice of annotations, the sets of variants being tested can be very different from each other, which would have downstream consequences on the observed results.

Of the variant prioritization methods we implemented in this study, we achieved the greatest power when using tissue-specific TURF scores. Using a combined-annotation scoring system, such as TURF, comes with advantages compared to using individual annotations. For one, it minimizes the multiple-testing burden because multiple annotations are already built into the scoring system without having to test them individually. Additionally, combined-annotation scoring systems also help eliminate any bias introduced by the manual selection of annotations. Manual selection of annotations relies on the investigator’s pre-conceived notions of which genomic regions may or may not be relevant or functional, and therefore has the potential to introduce irrelevant data or miss sources of true signal. Previous investigators who have focused on a few specific genomic regions (e.g. promoters, UTRs) have themselves pointed out that not all possible classes of noncoding regulatory elements were considered in their study [12], which could allow for other important regions to be missed. Zhou et al. have provided further evidence of the utility of combined-annotation scoring systems in their work in which they detected a significant burden of mutations affecting transcriptional and post-transcriptional regulation in ASD probands as compared to their unaffected siblings, using their Disease Impact Score [41]. We note that some differences in results between our group’s work and that of Zhou’s could be attributed to the fact that their prioritization scoring method was trained using a set of mutations specifically associated with disease from the Human Gene Mutation Database, while TURF was trained on SNVs associated with general regulatory function.

Different methods for identifying dnSNVs will yield different lists of mutations, even when starting with the same sequencing data or variant calls. These differences in lists between research groups can negatively affect reproducibility, and in fact we provide evidence that when working with the intersection of dnSNVs from other groups we can improve the overall ability to detect associations. When comparing the sets of mutations identified by different groups, including our own, it is encouraging to see that many of the same dnSNVs can be reproduced across groups (Fig 1c). We can reasonably expect to have higher confidence that the intersections of the sets represent true dnSNVs. Indeed, the fact that the intersection of our dataset and Zhou et al. yielded a stronger enrichment for DIS scores than either dataset alone suggests that the intersection is itself enriched for regulatory variants as compared to variants in the disjoint set. This is consistent with the possibility that variants discovered by only one group may be more likely to be false positives. We do note that some differences in dnSNV sets may be attributed to the fact that some studies included families that others did not, and there was little consistency in dnSNV identification methods across studies. Taking this into account, we suggest that improved methods of dnSNV identification and validation are likely to generate substantial improvements.

We were surprised to note that the use of Wilcoxon rank-sum tests instead of FET had disparate effects when using TURF brain vs DIS scores. Specifically, TURF brain scores performed better with FET (FET p=0.5; Wilcoxon rank-sum p=0.89) whereas DIS performed better with Wilcoxon rank-sum (Wilcoxon rank-sum p=0.071; FET p=0.214). This suggests that different prioritization scores may include biases that affect our ability to detect associations with a phenotype of interest. For example, while dnSNVs scoring within the top-5% of TURF brain-specific scores are modestly (but not significantly) enriched in probands, proband dnSNVs do not systematically rank higher than sibling dnSNVs based on their Wilcoxon rank-sum test results. By contrast, the reverse is true for DIS. While it is not immediately clear what may be responsible for the difference, it raises the question of whether a single significant test result can be considered definitive evidence of correlation or whether hidden structure may sway test results if only a single testing method is used, or whether disparate test results reflect shortcomings in the quality of a given annotation. This suggests that it may behoove researchers to compare results across different testing methods, giving preference to annotations that show consistent performance regardless of method.

Taking all these findings into account, we can make several recommendations for testing associations between genetic disorders and rare *de novo* variants:

1. Start with a high-confidence set of dnSNV calls, possibly leveraging intersections with other published datasets.
2. Select annotations and/or training features relevant to the tissue and/or phenotype of interest. Our results showed that brain-specific TURF scores outperformed generic TURF by a wide margin. Likewise, DIS outperformed brain-specific TURF in average score rank in probands and vs siblings, the difference being that the DIS model was trained on disease-specific regulatory variants, not general regulatory variants.
3. Do not rely on intuition alone in selecting annotations. Currently available machine-learning models do a better job of isolating signal from noise among a large and varied set of individual annotations.
4. Improving prioritization scores rather than increasing sample size is more likely to yield positive results. In particular, choosing training data that is relevant to the tissue or phenotype under study is of critical importance.

## Supporting information

Table of relative risk

Annotated SNVs

## Data and code availability

There are restrictions to the availability of source data from the Simons Simplex Collection (SSC) due to their data access policies. Approved researchers can obtain the SSC population whole-genome dataset described in this study by applying at https://base.sfari.org/.

## Supplemental information

SSC_dnSNVs.hg19.annotations.bed.zip - Table of annotated dnSNVs SSC_dnSNV_FET_relRisk.xlsx - Table of FET results (with Relative Risk)

## Acknowledgments

We are grateful to all of the families at the participating Simons Simplex Collection sites, as well as the principal investigators (A. Beaudet, R. Bernier, J. Constantino, E. Cook, E. Fombonne, D. Geschwind, R. Goin-Kochel, E. Hanson, D. Grice, A. Klin, D. Ledbetter, C. Lord, C. Martin, D. Martin, R. Maxim, J. Miles, O. Ousley, K. Pelphrey, B. Peterson, J. Piggot, C. Saulnier, M. State, W. Stone, J. Sutcliffe, C. Walsh, Z. Warren, E. Wijsman).

We thank Stephanie Bielas for helpful discussions. We thank Anshika Srivastava for her help in applying for SSC data access. We thank Shengcheng Dong for her help in generating TURF annotations. We also want to thank all members of the Boyle Lab for their support and constructive feedback.

This project was supported by NIH U24 HG009293. CC was supported by the University of Michigan Rackham Merit Fellowship and the Training Program in Bioinformatics (T32GM070449).

## Declaration of interests

The authors declare no competing interests

## Notes

### Competing Interest Statement

The authors have declared no competing interest.

